# Evaluating Flavoprotein Fluorescence Imaging as a Biomarker of Early Retinal Ganglion Cell Mitochondrial Stress

**DOI:** 10.1101/2025.08.27.672228

**Authors:** Tiffany M. Heaster-Ford, Pooja Teotia, Tom Truong, Jeffrey W. Hofmann, Miriam Baca, Shawnta Y. Chaney, Justin Elstrott

## Abstract

**Purpose:** Retinal neurodegeneration is difficult to monitor due to insensitive disease endpoints. Mitochondrial dysfunction and oxidative stress are promising early biomarkers of retinal ganglion cell (RGC) degeneration. This study investigates dynamics of flavoprotein fluorescence (FPF), a non-invasive mitochondrial oxidative stress measure, and sensitivity to early neurodegeneration and neuroprotection *in vitro* and *in vivo*.

**Methods:** FPF activity in response to neurodegeneration and neuroprotection were characterized in vitro in wild-type (WT) and SARM1 knockout (SARMKO) human embryonic stem cell-derived RGCs with and without Vacor treatment over 24 hours and confirmed with mitochondrial reactive oxygen species (ROS) measures. Further FPF evaluation was explored *in vivo* using the optic nerve crush (ONC) model in WT and SARMKO mice to compare early RGC stress detection within rodent retinas.

**Results:** *In vitro* FPF intensities in WT RGCs increased within 8 hours of degeneration induction, preceding significant mitochondrial ROS production. Neuroprotective SARMKO RGCs maintained comparable FPF and ROS levels following insult. *In vivo* FPF changes were not observed in WT and SARMKO mice over 4 days following ONC, while only early retinal thickening was observed from OCT. Early FPF and OCT changes were not reflective of late RGC survival observed from *ex vivo* RGC soma and axon counts.

**Conclusions:** These findings highlight differences in FPF sensitivity to mitochondrial stress between simplified *in vitro* systems and complex *in vivo* rodent retinas. This study demonstrates the potential of FPF as an early neurodegeneration and neuroprotection endpoint *in vitro* while identifying limitations and areas of development for its translatability to preclinical *in vivo* assessment.

## Introduction

Retinal ganglion cells (RGCs) are the sole projection neurons connecting the retina to the brain, crucial for normal visual processing^1^. Their health is vital to prevent retinal neuropathies. Following visual stimulus detection by photoreceptors and signal propagation by bipolar cells, RGCs generate action potentials for further processing by interconnecting neurons and relay to the brain’s visual cortex^2,3^. Retinal insults like elevated intraocular pressure, ischemia, and trauma can progressively damage RGCs, leading to gradual vision loss^4,5^. RGC death and vision loss can occur slowly, challenging early detection for preventive intervention^6^. Intracellular mechanisms, including oxidative stress and mitochondrial dysfunction, precede RGC damage^7,8^, highlighting the need for sensitive biomarkers to detect early degeneration.

Mitochondria are crucial for RGC function, supporting their high metabolic demands by generating energy and regulating oxidative stress^9,10^. RGCs rely heavily on mitochondria for ATP production, enabling ion transport for action potential propagation^11,12^. Mitochondrial antioxidants neutralize reactive oxygen species (ROS) from the electron transport chain (ETC) and other cell stress responses, maintaining intracellular ROS and ion balance and preventing cell damage^13,14^. RGC injury disrupts mitochondria-driven ETC function and antioxidant maintenance, limiting energy supply and initiating mechanisms causing ROS and calcium accumulation and axon fragmentation^11,15^. Evaluating defects in mitochondrial health and ROS regulation could indicate early RGC degeneration and development of disease pathophysiology.

Flavoproteins, integral to the ETC, are indicators of mitochondrial health, acting as redox cyclers for ATP production^16,17^. Increased ROS due to mitochondrial dysfunction and imbalanced redox cycling skew flavoproteins toward an oxidized state^18,19^. Flavoprotein oxidation alters protein conformation, enabling blue light absorption and green autofluorescence emission^18,19^. This flavoprotein fluorescence (FPF) is elevated in high oxidative stress states linked to mitochondrial damage, highlighting its potential as an endogenous stress indicator. Retinal FPF imaging has been explored clinically for stratifying disease states across several retinopathies^20–26^. Preclinical FPF studies have focused on oxidative stress monitoring in RPE stress models, but characterization has yet to be extended to other retinal stress models, limiting understanding of its relevance and utility as a mitochondrial stress biomarker^27,28^. Clinically, cross-sectional studies have demonstrated a correlation between elevated FPF and increasing retinal disease severity, but have not explored longitudinal changes in FPF after RGC injury and protection^21,23,24^. Expanded preclinical assessment of FPF in an optic nerve injury model may offer valuable insights into its utility as a biomarker of early RGC stress in retinal neurodegenerative diseases.

This study investigates FPF dynamics in response to RGC stress during axon degeneration, examining *in vitro* and *in vivo* sensitivity to early defects following RGC injury. Neuroprotection effects on FPF changes were also explored. FPF response in both neurodegenerative and neuroprotective conditions was initially characterized *in vitro* with retinal monoculture models and validated against mitochondrial oxidative stress markers. FPF sensitivity was then assessed in *in vivo* optic nerve injury models for early detection of RGC stress and neuroprotection in intact rodent retinas. Overall, the study aimed to characterize FPF sensitivity to early RGC mitochondrial stress in both homogeneous *in vitro* cultures and complex *in vivo* environments to better understand its potential as an endpoint for early retinal neurodegeneration and neuroprotection.

## Methods

### Human ESC-RGC culture generation and degeneration model induction

Human RGCs (hRGCs) were generated from human embryonic stem cells (hESCs) using a small molecule directed differentiation protocol as mentioned previously^29^. Established RGC cultures were seeded at concentration of 4×10^5^ cells per well in 48-well glass-bottom imaging plates and maintained in Neurobasal medium (Gibco) supplemented with B27, forskolin (Sigma-Aldrich), and ciliary neurotrophic factor and given fresh media every 2 days. Cell cultures were incubated at 37℃ and 5% CO_2_ during establishment and imaging. RGC degeneration was induced by treatment with 35 μM Vacor, previously shown as a Sterile alpha and TIR motif containing 1 (SARM1) agonist that drives Wallerian axonal degeneration^30^. Vacor (N-(4-Nitrophenyl)-N′- [(pyridin-3-yl)methyl]urea) was synthesized in-house using a previously-established protocol^31^.

### Human RPE culture generation and oxidative stress model induction

Immortalized human RPE cells (ARPE-19; ATCC) were cultured in 96-well glass bottom plates at a concentration of 2.5×10^5^ cells per well and maintained in DMEM:F12 medium (Gibco) supplemented with fetal bovine serum (Gibco) and penicillin:streptomycin (Gibco). Intracellular oxidative stress was induced by 3 hour treatment with various cell oxidation agents and ETC inhibitors. The treatments applied were as follows: cell oxidizers – 25-100 μM hydrogen peroxide (H_2_O_2_) and 3-20 mM sodium iodate (NaIO_3_); ETC inhibitors – 10-250 μM Paraquat, 1-50 μM Rotenone, and 0.1-5 μM MitoParaquat.

### Animals

All animal handling and procedures were performed following review and approval of the Institutional Animal Care and Use Committee (IACUC) at Genentech and in compliance with all policies, guidelines, regulations established by IACUC and the National Institutes of Health. SARM1KO mice were purchased from Jackson Laboratory (B6.129 × 1-Sarm1tm1Aidi/J; RRID:IMSR_JAX:018069) and bred in-house. SARM1 wt/wt (WT) and SARM1 ko/ko (SARMKO) phenotypes were confirmed via genotyping for absence or presence of the reversed neomycin resistance gene at exons 3-6, respectively. All animals were housed with ad libitum access to food and water and subjected to 14h light/10h dark cycle.

### AAV-mediated mitochondrial labeling

AAV-Cox8a-mTagBFP2 was developed using mitochondria-specific COX8A targeting sequence fused to mTagBFP2 and packaged into AAV_PHP.eB_ vector serotype by VectorBuilder at a production titer of 1.0 x 10^13^ vg/mL. RGC cultures were infected with AAV_PHP.eB_-Cox8a-mTagBFP2 at a multiplicity of infection of 250 for at least 4 days prior to treatment to promote mitochondria-targeted TagBFP2 expression and visualize mitochondrial transport.

### In vitro imaging and analysis

Imaging of RGC and RPE cultures was performed on a Nikon Ti-E spinning disk confocal microscope at 60x and 40x magnification, respectively. Live-cell imaging was conducted in an environmental chamber for maintenance of standard temperature and atmospheric conditions. All RGC imaging endpoints were collected in control and Vacor-treated conditions at 4, 8, and 24 hours following treatment. RPE images were collected 3 hours post-treatment with all cell oxidizers and ETC inhibitors.

### Mitochondrial imaging and analysis

Mitochondrial mTagBFP2 expression in RGC cultures was used to monitor mitochondrial localization within RGC axons and cell bodies. Mitochondria in RPE cultures were identified by live staining with Biotracker 405 following manufacturer protocol. Fluorescence emission was detected with a 420-470 nm bandpass filter upon 405 nm excitation. Mitochondrial segmentation was performed by thresholding mitochondrial fluorescence images to obtain binary mask image series for segmentation and tracking via an previously established, open-source MATLAB-based algorithm^32^

### Mitochondrial ROS imaging and analysis

MitoSox Green or Red mitochondrial superoxide dyes (Invitrogen) were used to assess mitochondrial superoxide production as a marker of ROS levels. Working solutions of MitoSox Green and Red were prepared at a concentration of 1 μM or 500 nM, respectively, in Hank’s balanced salt solution according to manufacturer’s instructions. RGC and RPE cultures were stained with Mitosox Green and Red, respectively, 30 minutes prior to imaging. MitoSox signal and mTagBFP2/Biotracker fluorescence were simultaneously imaged to ensure mitochondrial specificity of MitoSox signal. MitoSox fluorescence was excited at 488 nm and filtered over 500-555 nm. MitoSox intensities were quantified by generating binary masks based on thresholded mTagBFP fluorescence applied to raw MitoSox images and assessing the mean MitoSox intensity from each masked fluorescence image.

### In vitro FPF imaging and analysis

Endogeneous FPF signal was measured as a label-free marker of mitochondrial oxidative stress in *in vitro* control and Vacor-treated RGC and cell oxidizer/ETC inhibitor-treated RPE cultures. Here, mTagBFP2 or Biotracker fluorescence was similarly used as a marker of mitochondrial signal specificity. FPF emission was collected between 500-555 nm following 488 nm illumination. Thresholded mTagBFP/Biotracker fluorescence masks were applied to raw FPF images, and mean FPF intensity was quantified for all masked fluorescence images to evaluate FPF changes.

### Optic nerve crush injury model induction

Unilateral optic nerve crush (ONC) was performed in the right eye for all animals, and the contralateral left eye remained uninjured to serve as a control. Prior to the surgery, animals were anesthetized by intraperitoneal injection of ketamine (70-80 mg/kg body weight) and xylazine (15 mg/kg body weight) and maintained under anesthesia on a warming pad for all procedures. Proparacaine hydrochloride 0.5% eye drops (Sandoz) was applied as a topical anesthetic. Then, a subcutaneous injection of 3.25 mg/kg Buprenorphine (Ethiqa XR; Fidelis Animal Health) was given as a post-operative analgesic. Using a surgical scope for guidance, a fine incision was made in the superior-temporal conjunctiva, and the optic nerve is exposed using Dumont #5 forceps (Fine Science Tools). The crush injury was performed by positioning fine-tipped, self-closing forceps (Fine Science Tools) 1-2 mm distal to the globe and applying constant pressure directly to the nerve for 10 seconds. Post-operative care included incision closure and application of a topical antibiotic ointment (Neomycin and Polymyxin B Sulfates and Bacitracin Zinc Ophthalmic Ointment, Bausch & Lomb). All animals were monitored for recovery from anesthesia on a warming pad until fully awake and ambulatory.

### In vivo retinal FPF and OCT imaging

Prior to imaging, animals were anesthetized using isofluorane in an induction chamber prior to transition into a specialized animal positioner equipped to maintain a steady plane of anesthesia and consistent body temperature throughout the entire imaging session. Tropicamide ophthalmic solution UPS 1% (Akorn) was first applied to both eyes to promote pupil dilation, followed by application of Systane lubricant eye gel (Alcon) for long-term hydration of the cornea during imaging. A 10-mm, #1.5 round glass coverslip or a specialized rodent contact lens were mounted over the imaging eye to extend the hydration effects and create optimal coupling of light transmission between the objective and the retina for imaging.

An upright epifluorescence-capable microscope (Bruker 2pPlus) was used to capture retinal FPF signal *in vivo.* Full-field excitation and emission of FPF was performed by triggering a 60 millisecond flash using a white light LED (ThorLabs) equipped with a 460/30 excitation, 495 nm long pass, and 520/40 nm emission filterset (Chroma) relayed through a 10x/0.5 long-working distance objective (Thorlabs). Images were acquired using a CCD camera (Olympus, model S97827) under control by ToupView camera software (Touptek). Mean retinal FPF intensities were quantified by averaging the per-pixel FPF intensities across the entire field of view for all images.

A custom-built wavefront sensorless adaptive optics retinal imaging system (Netra Systems, Inc.) equipped with spectral-domain OCT was used to acquire OCT volumes of retinal structure *in vivo*. A broadband light source with 851.3 nm center wavelength ± 112.5 nm was used for OCT imaging. Raster scanned volume images were acquired centered at the optic nerve head (ONH) in both eyes. Each OCT volume captured 2048x500x500 A-scans at a 100 kHz scan rate, corresponding to a 2 mm imaging depth. The laser source was collimated to achieve a 1.2 mm beam diameter at the cornea and power was limited to 1.4 mW. The system provided 1.5 μm axial resolution and 3 μm lateral resolution in tissue for OCT image acquisition. The thickness measurements from the retinal nerve fiber layer (RNFL) and ganglion cell complex (GCC) were obtained from recorded OCT volumes using previously established, open-source code for automated segmentation of volumetric OCT images in the rodent retina^33^.

### Flatmount preparation, immunofluorescence imaging, and analysis

Following completion of the imaging study, animals underwent isoflurane anesthesia followed by subsequent 1x PBS and 4% PFA perfusion for rapid fixation. Whole globes were excised and underwent a secondary fixation by immersion in 4% PFA for 1 hour prior to transfer and storage in 1x PBS at 4℃ until dissected. Retinal dissection from the globes was carefully executed and visualized with a stereomicroscope, and dissected retinas were maintained at 1x PBS at 4℃ prior to start of the staining protocol. Permeabilization and blocking of retinal tissue was performed by immersion of the retina in 1x PBSTC (0.3% Triton X-100, 0.1mM CaCl_2_) on an orbital shaker for 1 hour and transfer to 1x PBSTC + 10% normal donkey serum (NDS) on an orbital shaker for 2 hours. Following permeabilization and blocking, retinal tissues were then stained with anti-RBPMS primary antibody (1:500, Novus) in 1x PBSTC + NDS for 4 days and Alexa-647 (1:1000) secondary antibody in 1x PBSTC + NDS overnight, incubated on an orbital shaker at 4℃ and with multiple PBSTC washes between primary and secondary staining. Stained retinas were then mounted on glass slides and diagonal incisions were made radially around the ONH, followed by addition of ProLong Gold Antifade mounting media (Invitrogen) and coverslip application. Fluorescence images of RBPMS-Alexa647 signal across all flatmounts were collected using a Leica Thunder widefield fluorescence imaging system acquired at 20x magnification with excitation and emission at 640 nm and 662-738 nm, respectively. RBPMS+ cell counts were quantified by segmenting fluorescent cells using CellPose 3.0 and importing the segmented regions of interest (ROIs) into FIJI for ROI count quantification^34,35^. Briefly, a 6.5 mm^2^ ring centered around the ONH was created, then four 0.16 mm^2^ square ROIs were drawn at the edge of the ring in each section quadrant. ROIs contained within each box were quantified using the Object Inspector plugin in FIJI.

### Optic nerve section preparation, imaging, and analysis

Excised rodent optic nerves were fixed for at least 24 hours immersed in modified Karnovsky’s fixative (2% paraformaldehyde and 2.5% glutaraldehyde in 0.1 M sodium cacodylate buffer, pH 7.2, Electron Microscopy Sciences, Hatfield, PA, USA) at 4℃. Nerves were washed in 0.1M Sorensen’s buffer (Electron Microscopy Sciences) and post-fixed with agitation in 1% osmium tetroxide (Electron Microscopy Sciences) in 0.1M Sorensen’s buffer for 24 hours at 4℃. Nerves were washed in 0.1M Sorensen’s buffer and incubated en bloc in 2% (w/v) uranyl acetate at room temperature for an hour. A final wash in 0.1M Sorensen’s buffer was performed before nerves were dehydrated with agitation through a sequence of 70, 95, and 100% ethanol, twice each for 10 minutes and followed by propylene oxide twice for 10 minutes. Tissues were infiltrated with eponate 12 resin (Ted Pella, Redding, CA) with agitation as follows: 1:1 eponate/propylene oxide for 12 hours, followed by 2:1 eponate/propylene oxide for 4 hours followed by pure eponate for 4 hours. The nerves were embedded in eponate in flat silicone embedding molds and polymerized at 70℃ overnight. Blocks were sectioned on an EM UC7 automated ultramicrotome (Leica Biosystems) equipped with a DIATOME (Quakertown, PA, USA) diamond knife and set to cut 1 μm sections at a 6-degree knife angle at 1.60 mm/s. Sections were collected on Superfrost positively charged glass slides (Thermo Scientific, Waltham, MA, USA). Slides were stained with 2% P-phenylenediamine ([PPD]; Sigma Aldrich, St Louis, MO, USA) in 50% ethanol for 1 minute, rinsed in water for 10 minutes, air dried, and coverslipped with Sakura (Torrance, CA, USA) Tissue-Tek Glass mounting medium. Stained sections were imaged using an Olympus VS200 slide scanner under oil immersion at 40x magnification. Axon segmentation and counting were performed using the Axonet 2.0 software^36^.

### Statistical analysis

All data are represented as mean ± standard error of the mean (SEM). Graphical representations of data and statistical analysis were generated using GraphPad v10.4.1. Comparisons between two-sample data were analyzed using unpaired t-tests, and multiple comparisons analysis was performed using either using ordinary or Welch’s one-way ANOVA with Tukey’s or Dunnett’s T3 multiple comparisons testing or two-way ANOVA with Sidak’s multiple comparison testing, denoted in the respective figure legend. Linear regression analysis was used for all correlative assessments between imaging and histological endpoints. Statistical significance was p<0.05 for all data.

## Results

### FPF is elevated in response to retinal stress and neurodegeneration in vitro

*In vitro* RGC degeneration was modeled using WT and SARMKO hESC-derived RGC cultures treated with Vacor, and FPF and MitoSox intensities were periodically measured over 24 hours in untreated and Vacor-treated cultures to assess and confirm mitochondrial stress preceding degeneration (Fig 1A). FPF images from WT RGCs demonstrated qualitative differences in FPF signal before and after Vacor-induced RGC degeneration (top row, Fig 1B). In contrast, SARMKO RGCs did not display pronounced differences in FPF signal in the presence or absence of Vacor treatment, suggesting that FPF may reflect mitochondrial dysfunction resulting from SARM1 activation (bottom row, Fig 1B). Quantitative comparison of FPF intensities revealed a gradual FPF signal increase in Vacor-treated WT RGCs over the acute timecourse, yielding significantly higher FPF intensities than untreated RGCs at 8 and 24 hours post-insult (Fig 1C). Conversely, SARM1 inhibition in RGCs ameliorated FPF elevation in response to Vacor treatment throughout the timeframe of exposure, further supporting that FPF shows early sensitivity to RGC mitochondrial health (Fig 1D). The proposed relationship between FPF signal and mitochondrial ROS production was thoroughly characterized in a separate human RPE cell stress model correlating changes in FPF with changes of mitochondrial ROS in response to cell oxidation and ETC activity perturbations (Supplemental Fig S1A), which confirmed a robust correlation between mitochondrial ROS production and FPF levels (Supplemental Fig S1B). To further explore potential sensitivity differences between FPF and mitochondrial ROS levels, WT and SARMKO RGCs were stained with mitochondrial superoxide indicator MitoSox Green (Supplemental Fig S2) and quantified for intensity changes with or without Vacor exposure over a 24 hour timecourse (Fig 1E-F). Vacor increased MitoSox intensity over control WT RGCs solely at 24 hours post-treatment (Fig 1E). Despite MitoSox signal’s dependence on upregulated oxidative phosphorylation^37,38^, control and Vacor-treated SARMKO RGCs did not exhibit significantly different MitoSox intensities at any timepoint (Fig 1F). These findings suggest FPF may offer earlier sensitivity to mitochondrial stress associated with neurodegeneration.

**Figure 1.**
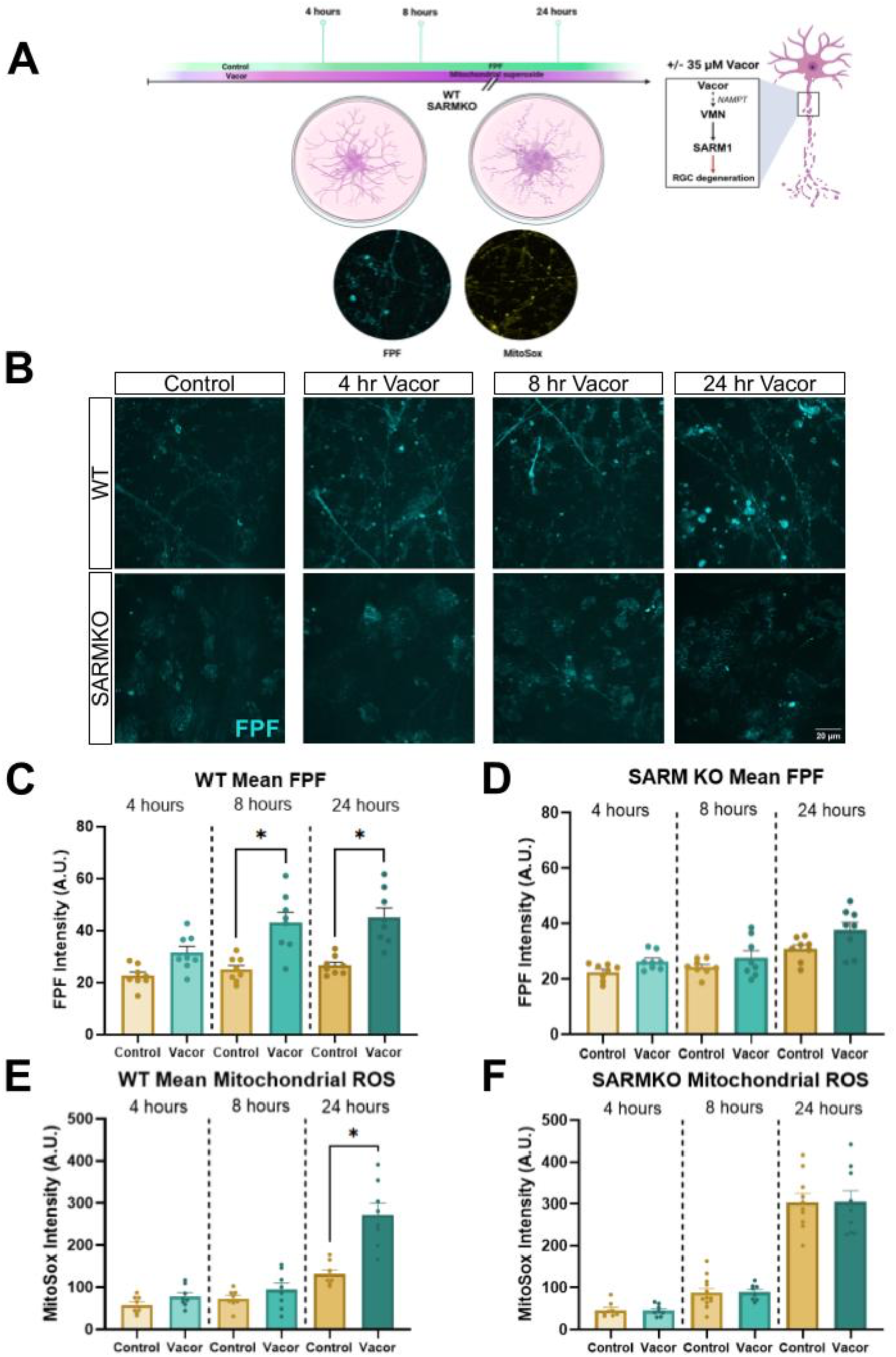
*In vitro* evaluation of acute FPF response after inducing RGC degeneration. **A.** Setup for FPF assessment in the *in vitro* RGC degeneration model. WT and SARMKO hESC derived-RGC cultures were treated with 35 μM Vacor to induce RGC degeneration. FPF & mitochondrial superoxide via MitoSox Green fluorescence were measured at 4, 8, & 24 hours post-treatment. **B.** Representative FPF images demonstrated elevated FPF signal in Vacor-treated WT RGCs compared with control, while FPF in SARMKO RGCs displayed comparable signal between control and treated cells at all timepoints. Scale bar: 20 μm. **C.** Vacor-treated WT RGCs exhibited a progressive increase in mitochondria-masked FPF signal over time and was significantly elevated above control RGCs at 8 hours and 24 hours post-insult. **D.** FPF in SARMKO RGCs was not significantly different between control and Vacor treatment conditions at any time point. **E.** Vacor yielded an increase in mean mitochondrial-masked MitoSox Green intensity over control WT RGCs solely at 24 hours post-treatment. **F.** No significant differences in MitoSox intensity were observed between control and Vacor-treated SARMKO RGCs. All data are represented as mean±SEM and evaluated using Welch’s one-way ANOVA with Dunnett’s T3 multiple comparisons tests. *p<0.05. N = 8 FOVs, 3 replicates/condition.

### In vivo FPF does not change significantly in response to early neurodegeneration following rodent ONC injury

We next tested whether the early, robust FPF increase observed *in vitro* after SARM1 activation would translate to an *in vivo* model of optic nerve injury, and whether SARM1 inhibition would mitigate any observed FPF elevation. Previous *in vivo* optic nerve injury studies found that SARM1 inhibition reduced axon degeneration distal to the lesion site but did not prevent somal or proximal axon loss, which are primarily driven by signaling upstream of SARM1^39,40^. However, SARM1 inhibition protects RGC somas in less severe chronic intraocular hypertension models^41,42^, suggesting SARM1 may play a role in somal degeneration in milder or possibly earlier stages of optic nerve injury. We thus measured retinal FPF changes in the days immediately following optic nerve crush to see if early FPF differences were detectable between SARM WT and KO mice prior to and concurrent with significant RGC loss, typically observed by 3 days post crush^43,44^. *In vivo* FPF response was evaluated longitudinally in rodent retinas with and without optic nerve crush induction, measured at one day prior to crush induction and at day 2, 3, and 4 following crush prior to takedown for ex vivo histological correlation (Fig 2A). Representative FPF images suggested early FPF decline in both uncrushed and crushed WT eyes, followed by a modest intensity increase exclusively in crushed eyes at day 4 for both SARM WT and KO mice (Fig 2B-C). Quantification of FPF intensities in WT mice trended towards modest elevation of baseline-normalized FPF intensity measurements at day 4 post-crush but were not significant, demonstrating comparable FPF to uncrushed eyes overall (Fig 2D). Similarly, FPF intensities were relatively stable for crushed eyes in SARMKO mice and not significantly different from uncrushed eyes (Fig 2E).

**Figure 2.**
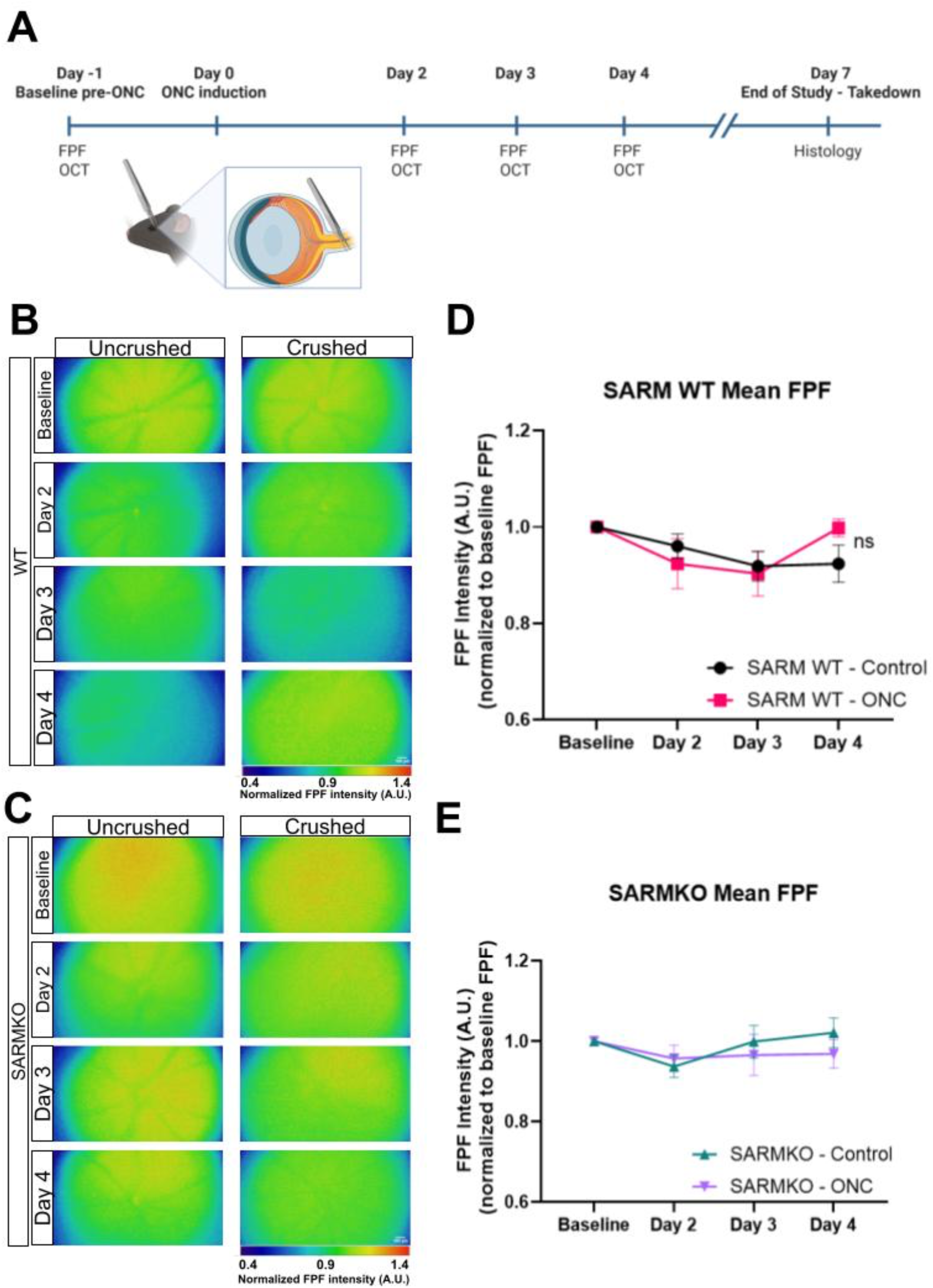
*In vivo* assessment of FPF sensitivity to neurodegeneration and neuroprotection following ONC. **A.** Study design for *in vivo* FPF assessment in the rodent ONC model. WT and SARMKO mice received unilateral ONC to cause progressive optic nerve damage. FPF was measured in intact and crushed eyes in all mice 1 day prior to ONC induction and at day 2, 3, & 4 post-crush. OCT volumes were also collected at each timepoint to observe concurrent changes in inner retinal layer integrity. Mice were taken down at day 7 to harvest retinas for histological analysis of RGC and axon survival. **B**. Representative FPF images in WT rodent eyes demonstrate non-significant changes in retinal fluorescence patterns with and without ONC induction over 4 days. Scale bar: 100 μm. **C.** Representative FPF images in uncrushed and crushed SARMKO rodent eyes also demonstrate comparable FPF levels over time. Scale bar: 100 μm. **D**. FPF response to ONC showed a trend towards mildly elevated normalized FPF intensity by day 4 post-crush, but overall FPF intensities were not significantly different from control. **E.** Normalized FPF intensities are similar between uncrushed and crushed eyes in SARMKO mice. All data are represented as mean±SEM and evaluated using two-way ANOVA with Sidak’s multiple comparison tests. N = 7-8 eyes per group.

### Early stages of degeneration following ONC similarly yield limited retinal structure changes

Structural OCT imaging was also performed at all study timepoints to evaluate structural differences as result of early neurodegeneration within the retina (Fig 2A). *En face* and cross-sectional OCT images were collected from WT (Fig 3A, left column) and SARMKO (Fig 3A, right column), displaying mild changes in retinal nerve fiber density and retinal layer architecture. Segmentation of the inner retinal layers from the inner limiting membrane (ILM) to the inner plexiform layer (IPL) enabled measurement of RNFL and GCC (ILM+RNFL/GCL+IPL) thicknesses corresponding to integrity of RGC axons and cell bodies post-optic nerve injury. Both RNFL and GCC thickness measures remained relatively stable over time in uncrushed WT and SARMKO eyes (Fig 3B-E). No significant change was observed in RNFL thicknesses between uncrushed and crushed eyes in WT and SARMKO mice (Fig 3B-C). GCC thickness was significantly greater in crushed eyes compared to uncrushed eyes in WT and SARMKO mice at Day 2 before gradually decreasing, suggesting early swelling in response to optic nerve injury prior to discernable structural loss (Fig 3D-E). Correlations of FPF intensity and retinal layer thickness changes from their respective baseline measurements revealed a weak relationship between FPF levels and both RNFL and GCC thickness (Supplemental Fig S3A-B), suggesting early FPF and OCT thickness changes after ONC may be mildly associated.

**Figure 3.**
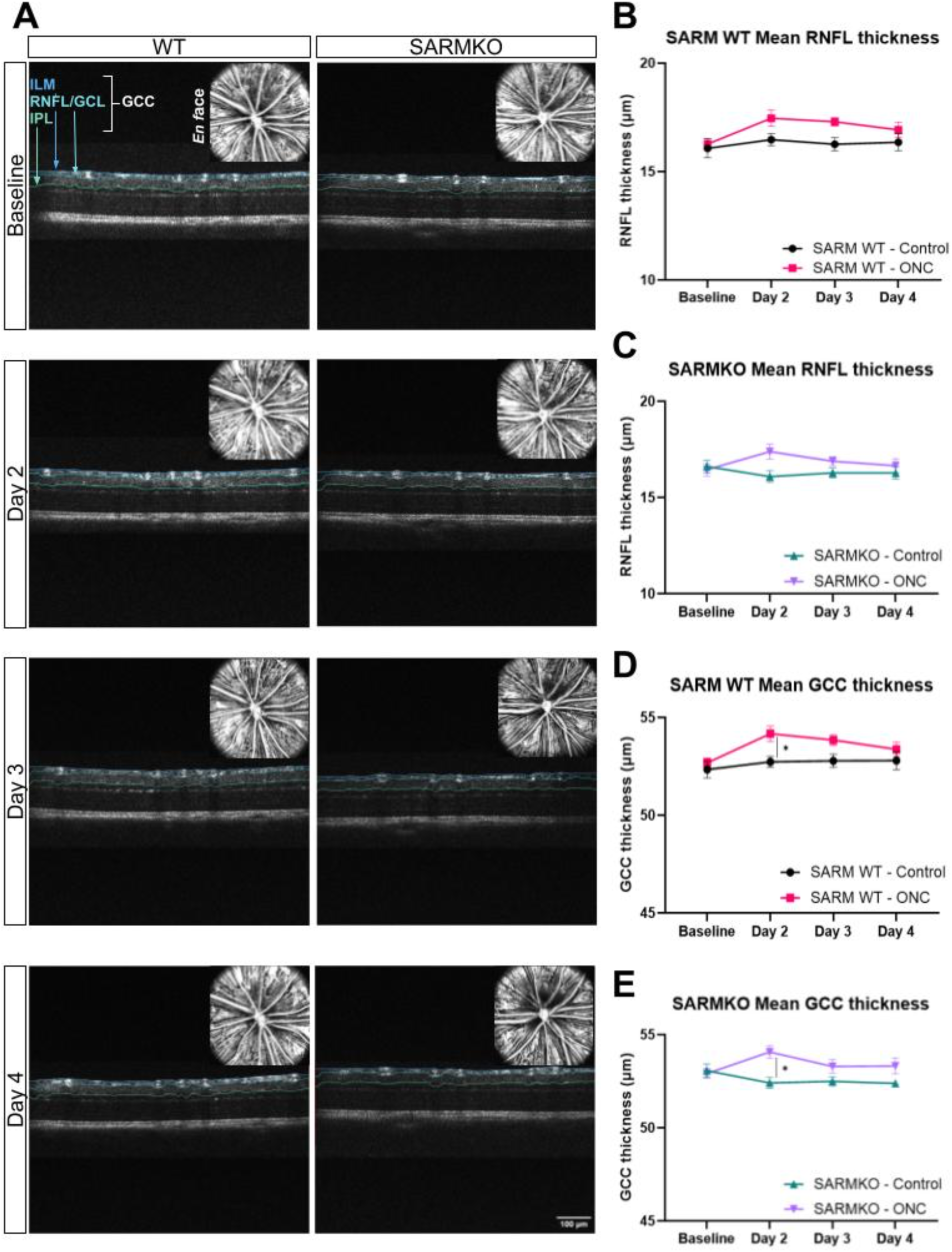
OCT retinal structure characterization during early degeneration following ONC. **A**. Representative images of cross-sectional and *en face* (inset) OCT in WT and SARMKO eyes visualize inner retinal layer and nerve fiber bundle thickness and organization following ONC. Scale bar: 100 μm (ILM – inner limiting membrane, GCL – ganglion cell layer, IPL inner plexiform layer). **B-C**. RNFL thickness measurements remained relatively stable over 4 days following ONC with minimal differences observed in uncrushed and crushed eyes in both **(B)** WT and **(C)** SARMKO mice. **D-E.** Crushed eyes in **(D)** WT and **(E)** SARMKO mice exhibited a modest increase in GCC thickness over uncrushed eyes solely at day 2 demonstrating early presentation and resolution of swelling. All data are represented as mean±SEM and evaluated using two-way ANOVA with Sidak’s multiple comparison tests. *p<0.05. N = 7-8 eyes per group.

### Early metabolic and structural changes do not reflect later optic nerve axon or retinal RGC survival following ONC

Standard histological measures evaluating cell-level changes in response to ONC injury were terminally assessed at Day 7, at which approximately 60-70% loss of RGC somas and axons is typically observed^43^. Segmented axons within PPD-stained optic nerve sections were quantified to determine differences in axon damage and loss between uncrushed and crushed WT and SARMKO optic nerves. Crushed nerves from both WT and SARMKO mice displayed a notable reduction in identified axons compared with uncrushed nerves (Fig 4A). However, more axons were preserved in the SARMKO crushed nerves compared to WT nerves (Fig 4A). Quantification of axon density confirmed WT and SARMKO axon loss in response to ONC, though higher axon density was observed in SARMKO nerves compared to WT nerves indicating axon protection (Fig 4B). Regression analysis of axon densities 7 days post-crush and change of FPF and OCT endpoints by day 4 post-crush was used to investigate whether early *in vivo* imaging metrics reflected late axon damage observed in *ex vivo* optic nerves. However, no significant correlations were observed between early changes in FPF intensity or OCT thicknesses and terminal axon density measures (Fig 4C-E).

**Figure 4.**
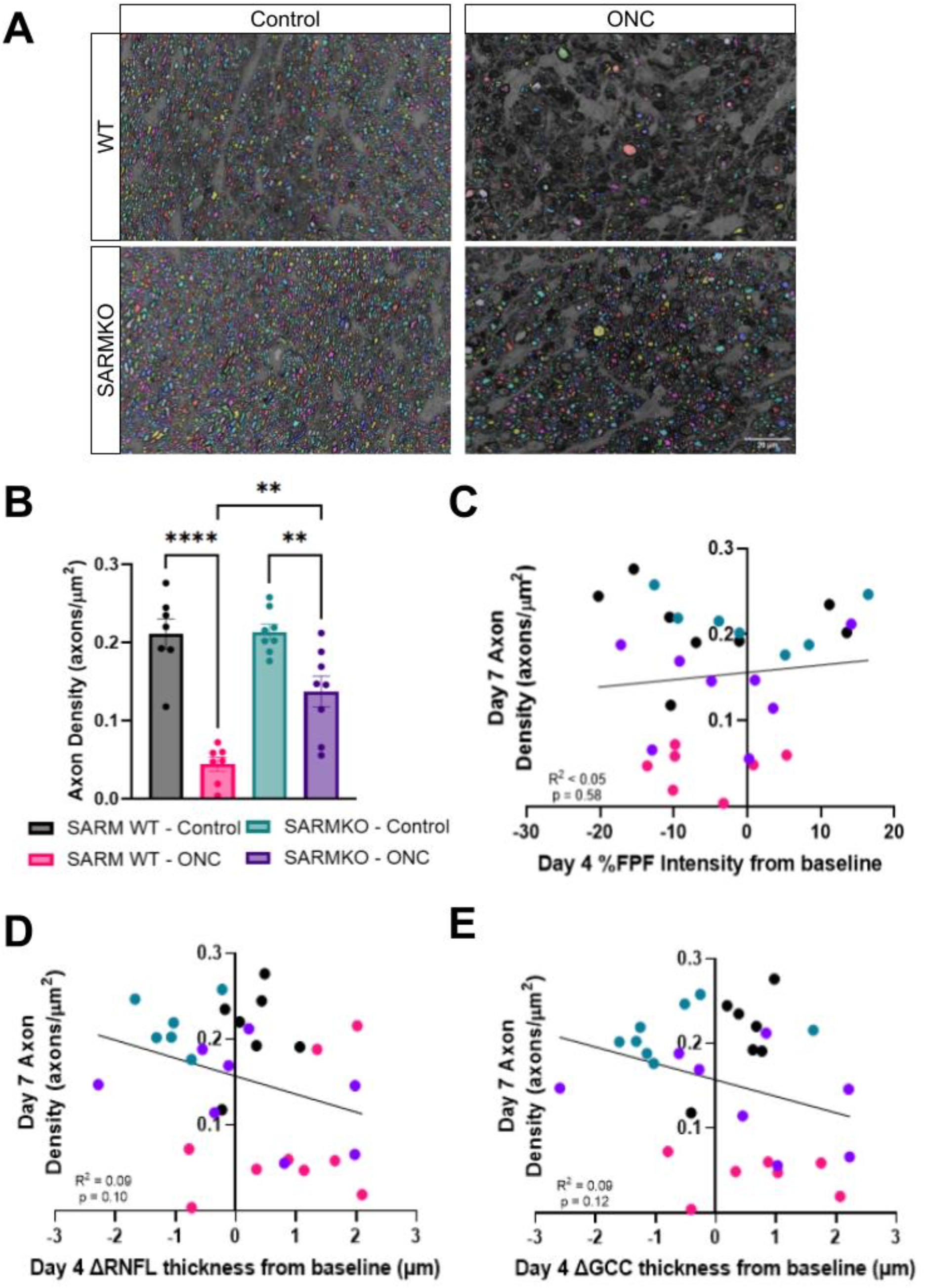
Histological endpoints of axon degeneration are not reflected by early *in vivo* imaging endpoints. **A.** Representative images of segmented axons from PPD-stained optic nerves sections from WT and SARMKO with and without crush induction demonstrate notable axon loss following ONC. Scale bar: 20 μm. **B.** Axon density quantification demonstrated significant axon loss in all crushed nerves by 7 days post-crush, and higher axon densities observed in SARMKO compared to WT crushed eyes demonstrating axon preservation. **C-E.** No significant correlations were observed between day 7 terminal axon density measures and day 4 changes in **(C)** FPF intensity, **(D)** RNFL thickness, and **(E)** GCC thickness from respective baseline measurements. Data in bar charts are represented as mean±SEM and evaluated using one-way ANOVA with Tukey’s multiple comparison tests. Linear regression analysis was used to evaluate correlation in all scatter plots. **p<0.01, ****p<0.0001. N = 7-8 eyes per group.

Previous work has demonstrated that inhibition of SARM1 activity protects RGC axons but not somas 14 days after ONC^42^. To confirm these findings, RBPMS+ RGC somas were segmented and counted to evaluate RGC degeneration within the retina in WT and SARMKO mice post-crush. Flatmounts of RBPMS-stained retinas displayed a dense field of RBPMS+ cells in uncrushed WT and SARMKO retinas (Fig 5A, left column). RBPMS+ staining in both WT and SARMKO retinas revealed lower density stain patterns, reflecting intermittent dropout of RBPMS+ cells (Fig 5A, right column). RBPMS+ cells were quantified from four 0.16 mm^2^ ROIs at the border of a central area (∼6.5 mm^2^) around the optic nerve head to provide representative sampling given density variations across the retina (Fig 5B). RGC soma counts confirmed significant RGC loss following crush induction, showing expected RGC loss in crushed eyes vs. uncrushed eyes for both WT and SARMKO 7 days post-crush (Fig 5C). As observed with axon density measures, there were no significant correlations between early changes in FPF intensity or OCT thickness and later terminal RGC counts (Fig 5D-F).

**Figure 5.**
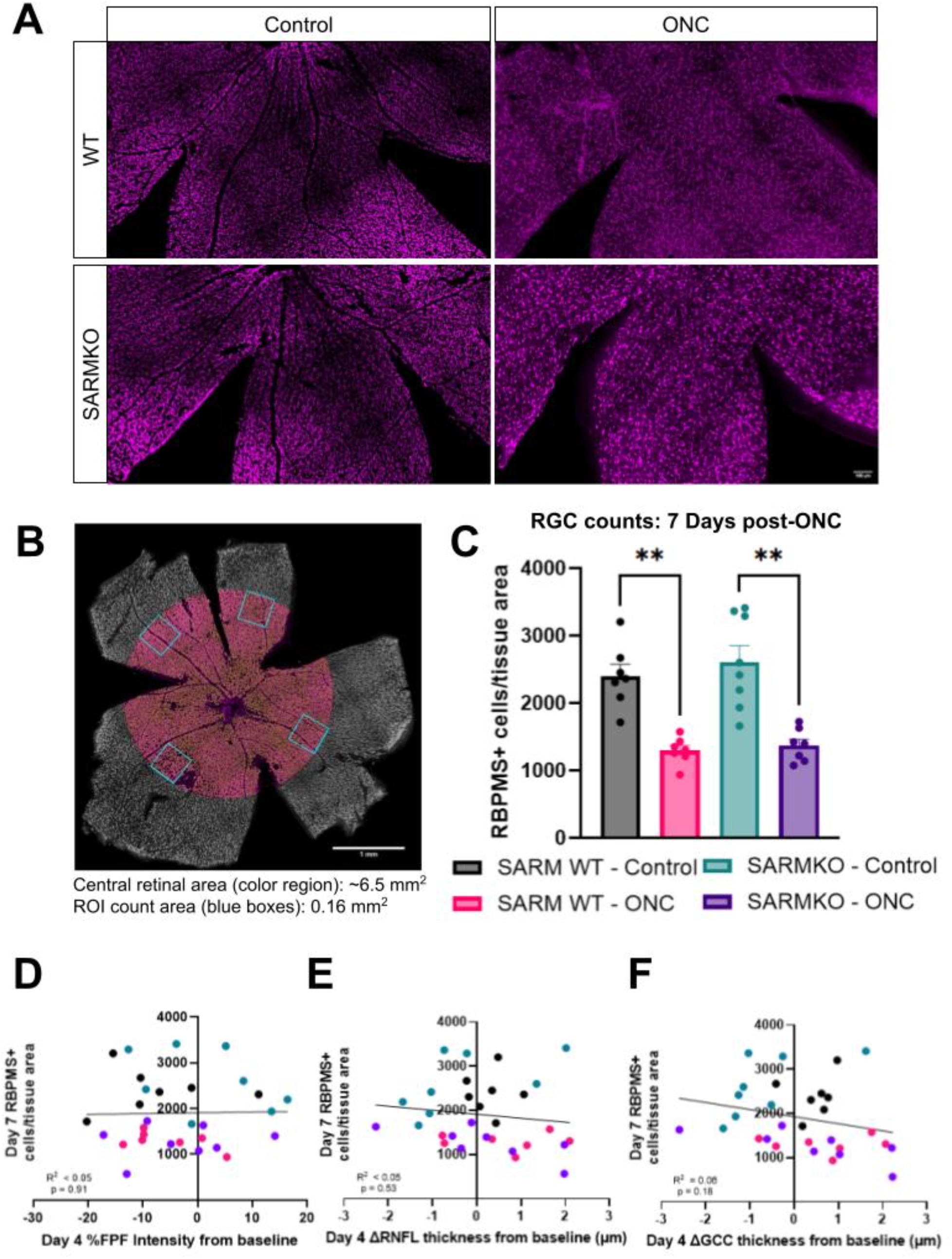
Histological endpoints of RGC loss are not reflected by early *in vivo* imaging endpoints. **A.** Representative images of RBPMS staining in flatmounts from uncrushed and crushed WT and SARMKO retinas display reduced RGC density following ONC. Scale bar: 100 μm. **B.** Diagram representing quantified areas of RBPMS+ RGC counts. A central area around the ONH measuring 6.5 mm^2^ was identified as a measurement boundary, and counts of positively-stained, segmented cells were measured from a 0.16 mm^2^ ROI along the boundary edge in each quadrant. **C.** RBPMS+ RGC counts were significantly lower in crushed eyes in both WT and SARMKO mice. **D-F.** No significant correlations were observed between day 7 terminal RBPMS+ RGC counts and day 4 changes in **(D)** FPF intensity, **(E)** RNFL thickness, and **(F)** GCC thickness from respective baseline measurements. Data in bar charts are represented as mean±SEM and evaluated using Welch’s one-way ANOVA with Dunnett’s T3 multiple comparisons tests. Linear regression analysis was used to evaluate correlation in all scatter plots. **p<0.01. N = 7-8 eyes per group.

## Discussion

RGC damage and death are hallmark features of several inner retinal neuropathies^45^. Retinal stressors, including increased intraocular pressure and ocular trauma, drive progressive RGC loss and vision decline^4,5^. Poor sensitivity of clinical endpoints to nascent retinal neurodegeneration limits the rate of early detection, which can delay diagnosis and treatment^46,47^. Mitochondrial and metabolic dysfunction present earlier than RGC somal or axon degradation, but current methods to monitor these early changes are largely restricted to cell and animal models, with limited translatability to clinical settings^7,8,48–50^. This underscores the need to explore novel, translatable biomarkers of oxidative stress associated with RGC degeneration.

This study investigated FPF, a non-invasive indicator of mitochondrial stress, as a potential biomarker of early RGC degeneration. We assessed FPF dynamics and sensitivity *in vitro* following direct activation of Wallerian degeneration in human RGC monocultures and *in vivo* in a rodent ONC model. *In vitro* FPF imaging detected time-dependent WT RGC defects preceding SARM1-activated axon degeneration not observed in SARMKO RGCs, confirming robust sensitivity in RGC cultures (Fig 1). These findings supported further development of FPF imaging for live, intact retinas, thus *in vivo* FPF validation was explored following ONC – a well-characterized model of acute RGC degeneration. Despite a trend towards FPF elevation, no significant changes in FPF were observed during early degeneration (4 days after ONC) (Fig 2). However, we did observe thickening of the RNFL and GCC 2 days after ONC in both SARMWT and KO retinas (Fig 3), suggesting that SARM1 inhibition does not affect early retinal changes such as retinal thickening or retinal cell loss associated with later timepoints following ONC^43,44^. Histological analysis 7 days after ONC confirmed RGC somal loss as well as axonal damage which was partially rescued in SARMKO animals as previously reported^42^, providing a benchmark to evaluate imaging sensitivity (Figs 4 and 5). However, the rapid progression of ONC-induced damage does not fully recapitulate the pathophysiology of chronic retinal neurodegenerative diseases, warranting further characterization of FPF and its potential utility as a disease progression biomarker in less severe, chronic ocular hypertension models.

Prior evaluation of FPF *in vitro* has provided early evidence of association with oxidative stress. These studies characterized oxidative stress-induced FPF in RPE cultures and *ex vivo* retinal tissues exposed to cell oxidizers or inflammatory stressors. We confirmed these reported relationships and evaluated pathway-dependent variations in human RPE cells treated with both mitochondrial complex inhibitors and cell oxidizers. FPF signal showed a strong correlation with increased ROS across treatment conditions, supporting its value as an indicator of retinal stress (Supplemental Fig S1). We then evaluated FPF directly in RGC monocultures to assess its relevance and sensitivity for RGC-specific pathologies. FPF elevation was detected 8 hours after Vacor-induced degeneration in WT RGCs, while SARMKO FPF levels remained unaffected after vacor treatment suggesting the FPF elevation observed in WT RGCs was driven by SARM1 activity (Figs 1C-D). Notably, mitochondrial superoxide levels in WT RGCs increased later compared to FPF levels (Fig. 1E), possibly due to rapid superoxide conversion to other ROS species (e.g., H_2_O_2_) not detectable by the indicator^47^. SARMKO RGCs showed delayed superoxide elevation without significant FPF changes (Figs 1D, F), potentially reflecting enhanced oxidative phosphorylation associated with SARM1 inhibition^37,38^. These findings underscore the utility of FPF for real-time, dynamic monitoring of RGC degeneration and neuroprotection *in vitro*.

Preclinical *in vivo* FPF evaluation has been limited in animal retinal disease models, highlighting a need for development to evaluate its utility as a translational biomarker of retinal stress. To date, preclinical *in vivo* retinal FPF has only been reported in healthy or aging, drusen-presenting rhesus macaques (Villafeurte-Trisonlini, et al. IOVS 2024;65:ARVO E-Abstract 2309; Ripolles-Garcia et al, IOVS 2025;66:ARVO E-Abstract 4766). To further characterize FPF dynamics in a rodent retinal neurodegeneration model, we developed a novel imaging setup to non-invasively monitor FPF in live rodent retinas prior to retinal insult and after degeneration was induced (Fig 2A). Longitudinal FPF assessment following ONC revealed minimal signal changes within 4 days (Figs 2D-E), contradicting our hypothesis that mitochondrial stress changes would manifest as early FPF elevation following ONC. This finding aligned with limited inner retinal thickness changes observed on OCT in this timeframe (Fig 3, Supplemental Fig S3), which preceded the more substantial retinal thinning and RGC loss typically observed at one week post-ONC^43,43,52^. To ensure successful model induction, neurodegenerative effects of ONC were confirmed with histological endpoints of RGC axon and somal density at day 7 post-crush (Figs 4B and 5C). We also confirmed SARM1-mediated neuroprotection of RGC axons (Fig 4B), but not somas (Fig 5C), as previously reported in the weeks following ONC^42^. The limited sensitivity of FPF and OCT in the early phase of degeneration was insufficient for correlation with these later histological changes (Figs 4C-E, 5D-F). These findings suggest that the complexity of the rodent retina may hinder FPF sensitivity *in vivo* during early degeneration.

This initial sensitivity characterization highlights current limitations and areas for improvement in FPF imaging for both *in vitro* and *in vivo* monitoring of retinal stress in cell culture and rodents, respectively. RGC monocultures in this study were maintained at physiologically relevant temperature and oxygen conditions, though we acknowledge that these systems do not fully recapitulate the retinal microenvironment. Further investigations additionally control of *in vitro* physiological parameters (e.g., pH, cell secretory factors) may better elucidate clear sources for the differences in FPF sensitivity *in vitro* and *in vivo*. Intrinsic ocular features and disease-induced changes have been shown to pose significant barriers to *in vivo* FPF signal collection. While our and previous *in vitro* mono- or co-culture studies have demonstrated up to a two-fold FPF increase in response to stressors^27,28^, this expected change in retinal FPF may be obscured *in vivo* due to fluorescence contribution from other tissues within the intact eye such as the cornea and lens. Clinical FPF studies also account for lens fluorescence and patient age, factors that can confound retinal FPF signals, but no equivalent methods exist for correcting rodent lens fluorescence^21^ (Muste et al. IOVS 2022;63:ARVO E-Abstract 219-F0066). Additionally, pathologies common to the ONC model and reported in diseased patients, such as edema and inner retinal thickening, have been suggested to obscure FPF signals and reduce sensitivity^43,44^ (Villafeurte-Trisonlini, et al. IOVS 2024;65:ARVO E-Abstract 2309; Pujari, et al. IOVS 2025;66:ARVO E-Abstract 3200). Advancing FPF imaging protocols and analyses to address these confounding factors will be crucial for improving its preclinical sensitivity and translational potential in detecting early retinal neurodegeneration and neuroprotection.

Early biomarkers of retinal neurodegeneration are essential for detecting retinal disease onset and progression and for guiding therapeutic development. This study provides initial preclinical evidence of the utility of FPF to monitor neurodegeneration and neuroprotection. We demonstrate FPF sensitivity to *in vitro* RGC mitochondrial stress and highlight its potential translatability for non-invasive assessment of early pathological changes in conditions linked to mitochondrial dysfunction. Further technological and experimental advancements are needed to establish FPF as a robust measure of retinal stress in rodent models and to bridge its clinical translatability. For example, correction for off-target fluorescence and consistent sampling from desired retinal regions adapted from FPF monitoring in higher order preclinical models and patients may improve the sensitivity of this measure in rodents and allow similar insight into global and regional changes gained from those applications. Feasibility of these enhancements for rodent FPF imaging and application in more physiologically relevant *in vitro* and *in vivo* models should be further explored. Overall, this work offers valuable insights into the capabilities and limitations of preclinical FPF imaging and may support improved clinical understanding of FPF for retinal stress monitoring in diseased patients.

## Acknowledgements

The authors would like to thank Taehoon Kim and Mitchell Geringer for guidance and insightful discussions on OCT imaging studies and *ex vivo* tissue analysis, respectively. The authors also would like to acknowledge the support provided by the necropsy team and laboratory animal resources for live animal and tissue collection management, Sarah Gierke and the Center for Advanced Light Microscopy for recommendations on live cell imaging, and Netra Systems Inc. for technical support for the adaptive optics retinal imaging system. Copy editing support was provided on 7/2/2025 using an internal Roche instance of OpenAI’s ChatGPT-4o.

