## Supplementary Material for "Evaluating Flavoprotein Fluorescence Imaging as a Biomarker of Early Retinal Ganglion Cell Mitochondrial Stress"

**Supplemental Figure S1. FPF signal correlates with mitochondrial ROS production in response to intracellular oxidative stress. A.** *In vitro* retinal oxidative stress was modeled by treating ARPE-19 cells with varied doses of cell-level oxidizers (hydrogen peroxide and sodium iodate) and mitochondrial complex inhibitors (rotenone, paraquat and MitoParaquat) for 3 hours. Changes in mitochondrial FPF signal relative to mitochondrial ROS levels were assessed by imaging mitochondria-specific fluorescence from FPF and MitoSox Red, a mitochondrial superoxide indicator. **B.** FPF intensity exhibited a strong, positive correlation with mitochondrial ROS production represented by MitoSox Red intensity in response to oxidative stress induced by both cell oxidation and ETC inhibition ($p<0.001$).

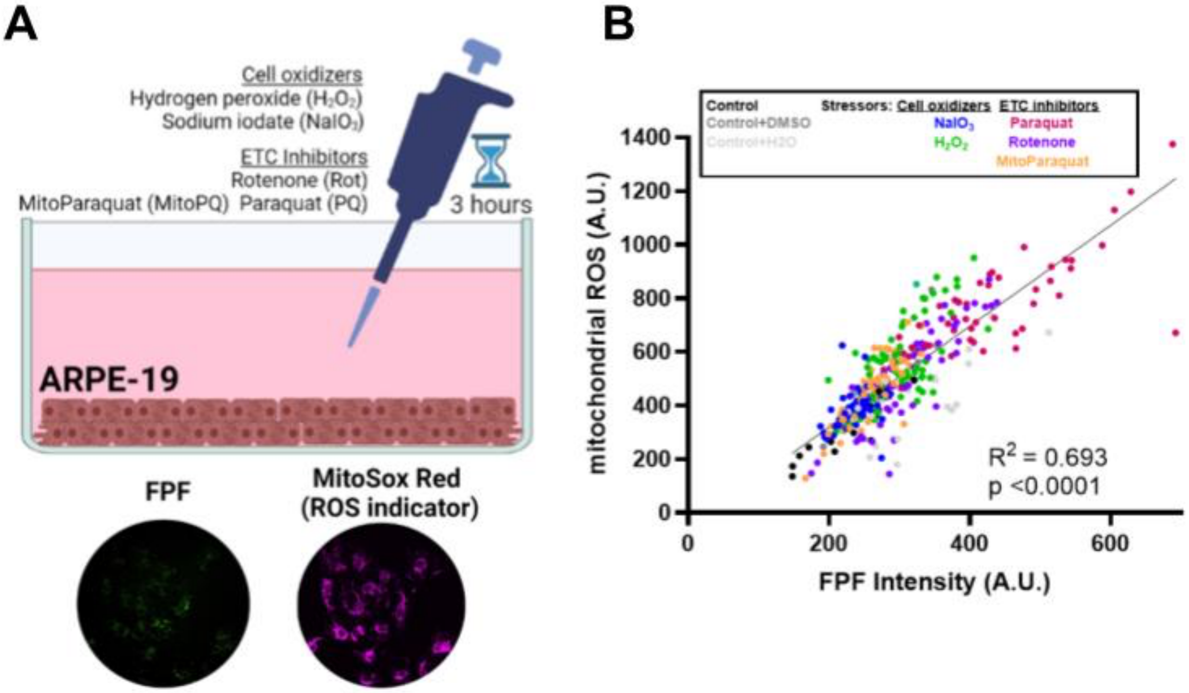

**Supplemental Figure S2. Representative MitoSox intensity images.** Representative images of WT and SARMKO RGCs with and without vacor treatment over 24 hours following staining with MitoSox Green, a mitochondrial superoxide indicator. Scale bar: 20 μm

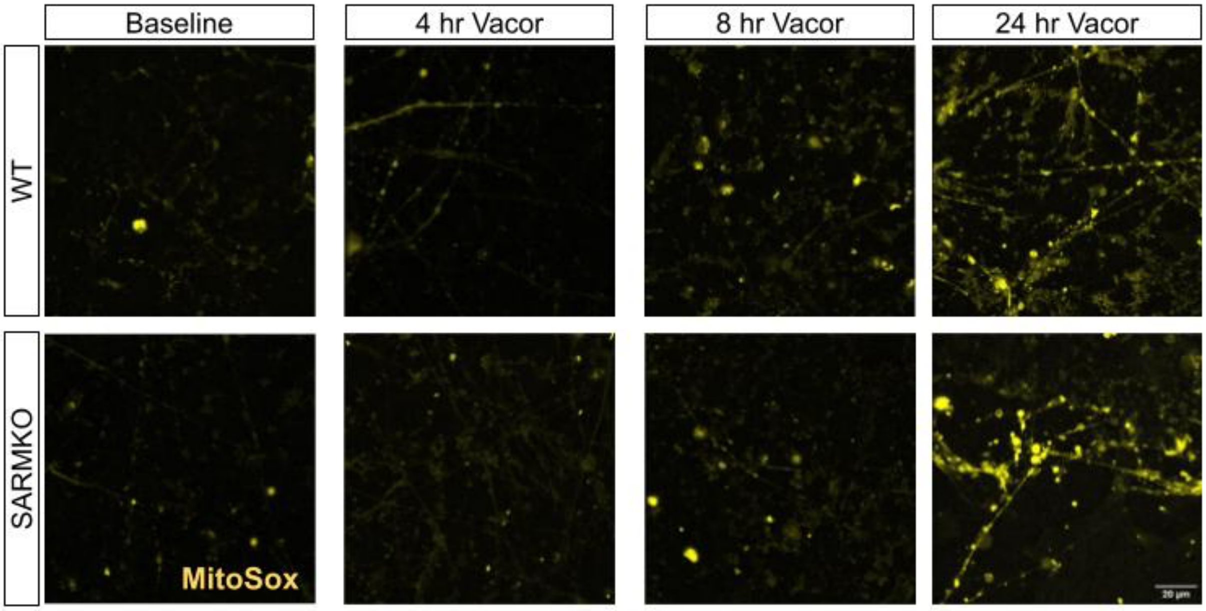

**Supplemental Figure S3. Change in FPF from baseline to Day 4 post-crush weakly correlates with change in RNFL and GCC thickness.** Linear regression analysis was performed to correlate changes in FPF intensity and OCT RNFL and GCC thickness between measurement at baseline (pre-crush) and Day 4 post-crush. **A-B.** FPF intensity changes were shown to be weakly correlated with **(A)** RNFL thickness (p = 0.015) and **(B)** GCC thickness (p = 0.016) changes.

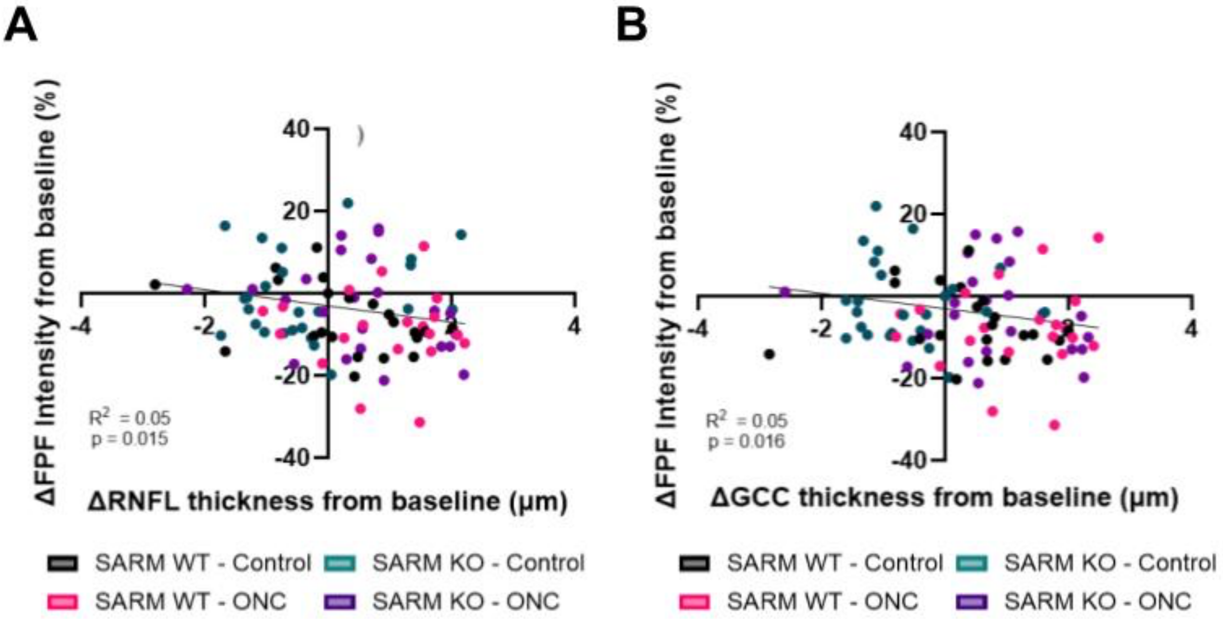
